# The combination of mitogenic stimulation and cell cycle arrest induces senescence in equine and human cartilage explants

**DOI:** 10.1101/2020.03.05.979872

**Authors:** Michaela E. Copp, Margaret C. Flanders, Rachel Gagliardi, Jessica M. Gilbertie, Garrett A. Sessions, Susanna Chubinskaya, Richard F. Loeser, Lauren V. Schnabel, Brian O. Diekman

**Author notes:** **Corresponding Author:** Brian Diekman, Thurston Arthritis Research Center, The University of North Carolina School of Medicine, 3300 Thurston Building, Campus Box #7280, Chapel Hill, NC 27599-7280. **Email addresses:** Michaela Copp Margaret Flanders Rachel Gagliardi Jessica Gilbertie Garrett Sessions Susanna Chubinskaya Richard Loeser Lauren Schnabel Brian Diekman.

## Abstract

**Objective:** Cellular senescence is a phenotypic state characterized by stable cell-cycle arrest, enhanced lysosomal activity, and the secretion of inflammatory molecules and matrix degrading enzymes. Senescence has been implicated in osteoarthritis (OA) pathophysiology; however, the mechanisms that drive senescence induction in cartilage and other joint tissues are unknown. While numerous physiological signals are capable of initiating the senescence phenotype, one emerging theme is that growth-arrested cells convert to senescence in response to sustained mitogenic stimulation. The goal of this study was to develop an *in vitro* articular cartilage explant model to investigate the mechanisms of senescence induction.

**Design:** This study utilized healthy articular cartilage derived from cadaveric equine stifles and human ankles. Explants were irradiated or treated with palbociclib to initiate cell cycle arrest, and mitogenic stimulation was provided by serum-containing medium (horse) and the inclusion of growth factors (human). The primary readout of senescence was a quantitative flow cytometry assay to detect senescence-associated β galactosidase activity (SA-β-gal).

**Results:** Irradiation of equine explants caused 25.39% of cells to express high levels of SA-β-gal, as compared to 3.82% in control explants (p=0.0031). For human cartilage, explants that received both mitogenic stimulation and cell cycle arrest showed increased rates of senescence induction as compared to baseline control (7.16% vs. 2.34% SA-β-gal high, p=0.0007).

**Conclusions:** Treatment of cartilage explants with mitogenic stimuli in the context of cell-cycle arrest reliably induces high levels of SA-β gal activity, which provides a physiologically relevant model system to investigate the mechanisms of senescence induction.

## Introduction

Osteoarthritis (OA) is a disease characterized by joint pain and progressive degradation of articular cartilage and other tissues of the joint^1,2^. As the most common chronic disease of the articular joint, OA produces a substantial burden on society and the economy^3,4^. Despite increasing knowledge about factors contributing to the progression of OA, there are no approved disease-modifying therapies^5^, leading to high rates of total joint replacement^6^. Risk factors for OA include obesity, joint injury, and genetic predisposition, with the most dominant risk factor being aging^7,8^. Cellular senescence has been described as a key phenotype associated with aging^9^, and there is mounting evidence that the accumulation of senescent cells in the joint during both aging and in response to injury contributes to the development of OA^10–14^. Senescent chondrocytes likely contribute to tissue degradation by producing pro-inflammatory and matrix-degrading molecules known collectively as the senescence-associated secretory phenotype (SASP)^15,16^. Significant advances have begun to unravel the role of senescence in OA and the therapeutic implications of such findings, including the potential for senolytic therapy as a potential disease-modifying therapy^13,17^. However, there has been less progress in understanding the underlying biologic processes that drive the accumulation of senescent cells in the joint space. With clinical trials that target senescent cells in the joint underway (e.g. NCT03513016), it is imperative to identify the physiological contexts that promote senescence induction^18^. The goal of this study was to investigate the mechanisms of senescence induction in articular cartilage through the use of explants from healthy equine and human cadaveric donors.

Senescent cells display stable cell cycle arrest even in an environment that would normally promote cell division, which distinguishes the senescent phenotype from quiescence^19^. Indeed, there is evidence that a pro-growth environment – termed expansion signals to account for stimuli associated with proliferation or cellular hypertrophy – drives senescence induction in cells harboring strong growth arrest (reviewed in ^20^). The upregulation of metabolic processes associated with an ongoing stress response, combined with absence of division, result in abnormally high lysosomal activity in senescent cells. This feature has been used to identify senescence through detection of β-galactosidase activity at the sub-optimal pH of 6.0 – the senescence-associated β-galactosidase (SA-β-gal) assay^21,22^. We recently used cartilage explants from p16^tdTomato^ knock-in senescence reporter mice^23^ to show that transforming growth factor beta (TGF-β1) and basic fibroblastic growth factor (bFGF), both of which are released from cartilage tissue in response to injury and degradation, were potent inducers of senescence by the measure of *p16*^*Ink4a*^ promoter activity^24^. In this study, we implement a quantitative flow-cytometry-based assay of SA-β-gal activity to demonstrate that the combination of cell-cycle arrest and cell-expansion stimuli induce senescence in equine and human cartilage.

## Materials and methods

### Acquisition of equine cartilage explants

Cartilage isolation was performed under IACUC approval at the North Carolina State University College of Veterinary Medicine from donor horses that were euthanized for reasons outside of this study. Horses were between 4 and 7 years of age and included 3 geldings and 4 non-parous mares. A series of 6 mm biopsy punches were taken from the femoral trochlear ridges of thoroughbred horses without known patellofemoral disease or any macroscopic signs of cartilage damage.

### Acquisition of human cartilage explants

Tali from cadaveric ankle joints were obtained from organ donors within 24 hours of death through a collaboration between Rush University Medical Center (Chicago, IL) and the Gift of Hope Organ and Tissue Donor Network (Itasca, IL) under a protocol approved by the Rush University institutional review board. Samples were shipped overnight with ice packs to the University of North Carolina at Chapel Hill. Donor samples were screened with a modified version of the 5-point Collins grading scale^25^ and only ankle joints with grades of 0 – 2 were used to avoid the confounding factor of cartilage degeneration. Explants of 5 mm were taken from the talar surface for culture. Tissue was used from a total of 19 donors – 13 males and 6 females ranging from 38 to 72 years of age. Donor information is included in a table alongside the appropriate figures.

### Explant culture for senescence induction

The experimental layout is provided as a schematic in Figure 1. Harvested equine explants were allowed to recover in 6-well plates for 3-7 days in the following control medium: DMEM/F12 medium (11330, Thermo Fisher Scientific, Waltham, MA), 10% fetal bovine serum (Seradigm 1500-500; VWR International, West Chester, PA, USA), penicillin and streptomycin (15140; Thermo Fisher Scientific), gentamicin (15750; Thermo Fisher Scientific), and amphotericin B (A2942; MilliporeSigma, Burlington, MA, USA). Half of the explants from each horse donor were irradiated with 10 Gy using a RS2000 Biological Irradiator, with the other half remaining as experimental controls. Post irradiation, the explants were cultured in the same control medium for 7 – 10 days before digestion for monolayer culture. Human cartilage explants were cultured in the same way as with equine explants, with the exception that mitogenic stimulation was included as an additional experimental factor. Mitogenic stimulus conditions were applied immediately after irradiation and consisted of control medium with the addition of 1 ng/mL TGF-β1 and 5 ng/mL basic fibroblastic growth factor (bFGF) (PHG9204 and PHG0264; Thermo Fisher Scientific). In additional experiments with human cartilage, palbociclib was used in place of irradiation as an additional mediator of cell cycle arrest. Palbociclib (PD-0332991 HCI, ChemShuttle, Wuxi, China) was applied after explant recovery and treatment lasted for 12 days at a concentration of 1 µM in complete media, with media being replaced every 2-3 days.

**Figure 1.**
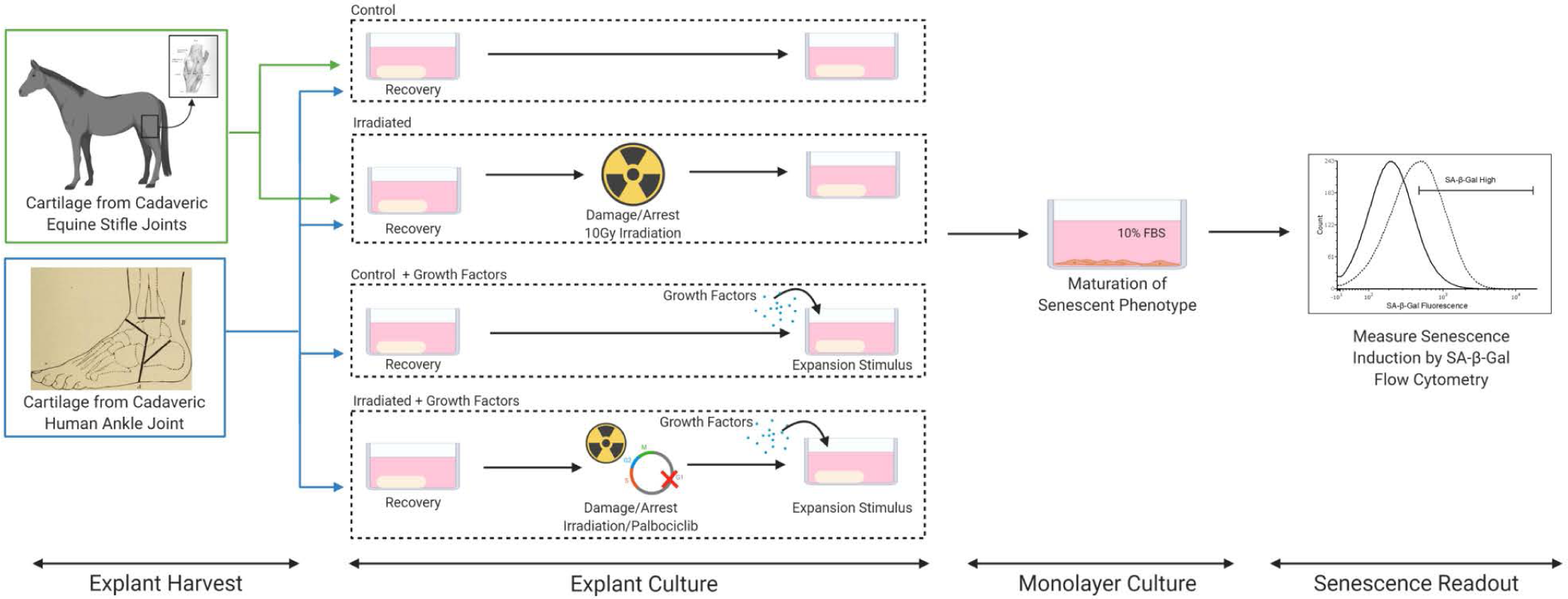
Experimental system schematic. Senescence was induced in healthy cadaveric cartilage by damaging explants with irradiation (10 Gy) or inducing cell cycle arrest with palbociclib (human explants only), culturing the explants in 10% FBS media (horse) spiked with growth factors – TGF-β1 and bFGF (human) – for 7 to 10 days, followed by enzymatic digestion and culture in monolayer for an additional 10 to 12 days. Senescence induction was assessed via SA- β-gal flow cytometry. Figure Created with BioRender.com.

### Monolayer culture of primary chondrocytes for maturation of senescent phenotype

Following senescence induction in explant culture, chondrocytes were isolated from cartilage by enzymatic digestion with Pronase (1 hour) and subsequently with Collagenase P (overnight) as described previously^26^. Isolated chondrocytes were resuspended in 1 mL of media and plated in 12-well plates at a concentration of 6.4×10^4^ cells per cm^2^. All cells (including those from the growth factor treated explants) were cultured in control medium, with media being changed every 2-3 days. The chondrocytes were cultured in monolayer for 10 – 12 days before SA-β-Gal flow cytometry analysis. Images of chondrocytes at the end of monolayer culture were taken with an EVOS FL microscope (Life Technologies, Carlsbad, CA).

### Flow cytometry analysis of SA-β-gal

SA-β-gal levels in the chondrocytes after senescence induction and monolayer culture were evaluated using the CellEvent™ Senescence Green Flow Cytometry Assay Kit (C10840; Thermo Fisher Scientific). The working solution was prepared by diluting the CellEvent™ Senescence Green Probe (1,000X) into the CellEvent™ Senescence Buffer that had been warmed to 37^°^C. Chondrocytes to be analyzed were washed twice with 1x DPBS (14190136, Gibco), incubated at 37^°^C for 5 minutes in Trypsin-EDTA (T4174; Sigma-Aldrich, St. Louis, MO), and neutralized with 50 µg/mL Soybean Trypsin Inhibitor (17075029, Thermo Fisher Scientific) with EDTA (46-034-CI, Corning, Corning, NY). The mixture was centrifuged at 800 xg for 6 minutes, washed with PBS, and fixed with 4% paraformaldehyde in PBS (from 16% stock, 43368, Alfa Aesar, Haverhill, MA) for 10 minutes at room temperature. Fixed cells were washed in 1% bovine serum albumin (A7906; Sigma Aldrich) in PBS and the chondrocytes resuspended in 100 µL of the working solution. The chondrocytes were incubated with the dye at 37^°^C for 2 hours at 300 rpm in a ThermoMixer C (Eppendorf, Hamburg, Germany). After incubation, the chondrocytes were washed with 1 mL of DPBS and resuspended in 200 µL of DPBS. The stained cells were filtered with a 30-µm strainer to achieve a single-cell suspension. Flow cytometry analysis was performed using an Attune NxT (Thermo Fisher Scientific) with a 488 nm laser, and analysis performed using FCS Express 8 software (De Novo Software, Glendale, CA, USA).

### Histology and Chondrocyte Cluster Analysis

Equine and human and cartilage explants at the end of the explant culture were fixed in 4% paraformaldehyde in PBS for 24 hours at 4^°^C and processed for paraffin embedding. Sections of 5 µm thickness were collected on Superfrost™ Plus slides (12-550-15; Thermo Fisher Scientific). Slides were stained with Hematoxylin (nuclei) and Eosin (cytoplasm) to detect cell cluster formation in explants. Representative bright field images were taken with an Olympus BX60 microscope. Chondrocyte cluster analysis was assessed from representative images of 2-3 independent explants from 4 different human donors. The number of chondrocytes in singlet, doublet, and triplet or more (triplet^+^) clusters were counted and quantified using the ImageJ imaging software (Fiji).

### Statistical analyses

Statistical analysis and plotting were performed using Prism 8 (GraphPad, La Jolla, CA, USA) and flow cytometry data was processed with FCS Express 8 (De Novo Software, Glendale, CA, USA). Data are plotted as individual points with the mean shown. The error bars indicate the 95% confidence interval (CI) and these values are noted in parentheses on the appropriate plots. Outliers were identified using Grubbs test with α = 0.05 and were excluded from subsequent analysis. Data from individual donors were excluded if the flow cytometry event count for any condition was less than 800 events. Outliers and excluded data are noted in the figure legends. Mean fluorescence intensity (MFI) data were normalized to the control condition for each donor to account for subtle differences in flow cytometry settings between days. Normalized MFI data in each treatment group were analyzed using a one-sample t-test against a hypothetical value of one. The percentage of SA-β-gal high cells was determined by introducing a cutoff at the mean plus two times the standard deviation of the control group. These percentage data were confirmed to be normally distributed by Shapiro-Wilk test and were analyzed using a paired Student’s t-test (equine) or Two-Way ANOVA with Tukey’s Multiple Comparisons Test (human).

## Results

### Induction of senescence in equine cartilage explants

Cartilage explants harvested from the equine stifle (patellofemoral joint) were induced to senescence in explant culture with irradiation followed by culture in monolayer for maturation of the senescent phenotype (overview of experimental design illustrated in **Fig. 1**). Chondrocytes from explants that had been irradiated revealed an enlarged and flattened morphology as compared to chondrocytes from control explants (**Fig. 2a**). Furthermore, the irradiated group contained a number of chondrocytes with lipid aggregates, which has been associated with senescence in other cell types^27^. Flow cytometry for SA-β-gal was used as a measure to quantify the induction of senescence in the cartilage explants. A representative SA-β-gal flow plot from one equine donor is included, showing the shift in SA-β-gal fluorescence between chondrocytes cultured from the control and irradiated equine explants (**Fig. 2b**). The region two standard deviations above the control mean fluorescence intensity (MFI) was delineated on the SA-β-gal flow plot and was used in this study to indicate the population of SA-β-gal high cells. Chondrocytes irradiated in explant culture had significantly increased SA-β-gal expression as compared to chondrocytes from control explants (2.40-fold increase, CI 0.22 to 2.58; p = 0.027 by one-sample t-test compared to a hypothetical value of 1; **Fig. 2c**). The percentage of cells with high SA-β-gal activity also significantly increased in the irradiated condition as compared to control (mean difference: 21.57%, CI 10.48 to 32.65%; p = 0.0031 by paired t-test; **Fig. 2d**). A table listing the age and sex of the equine donors is provided (**Fig. 2e**).

**Figure 2.**
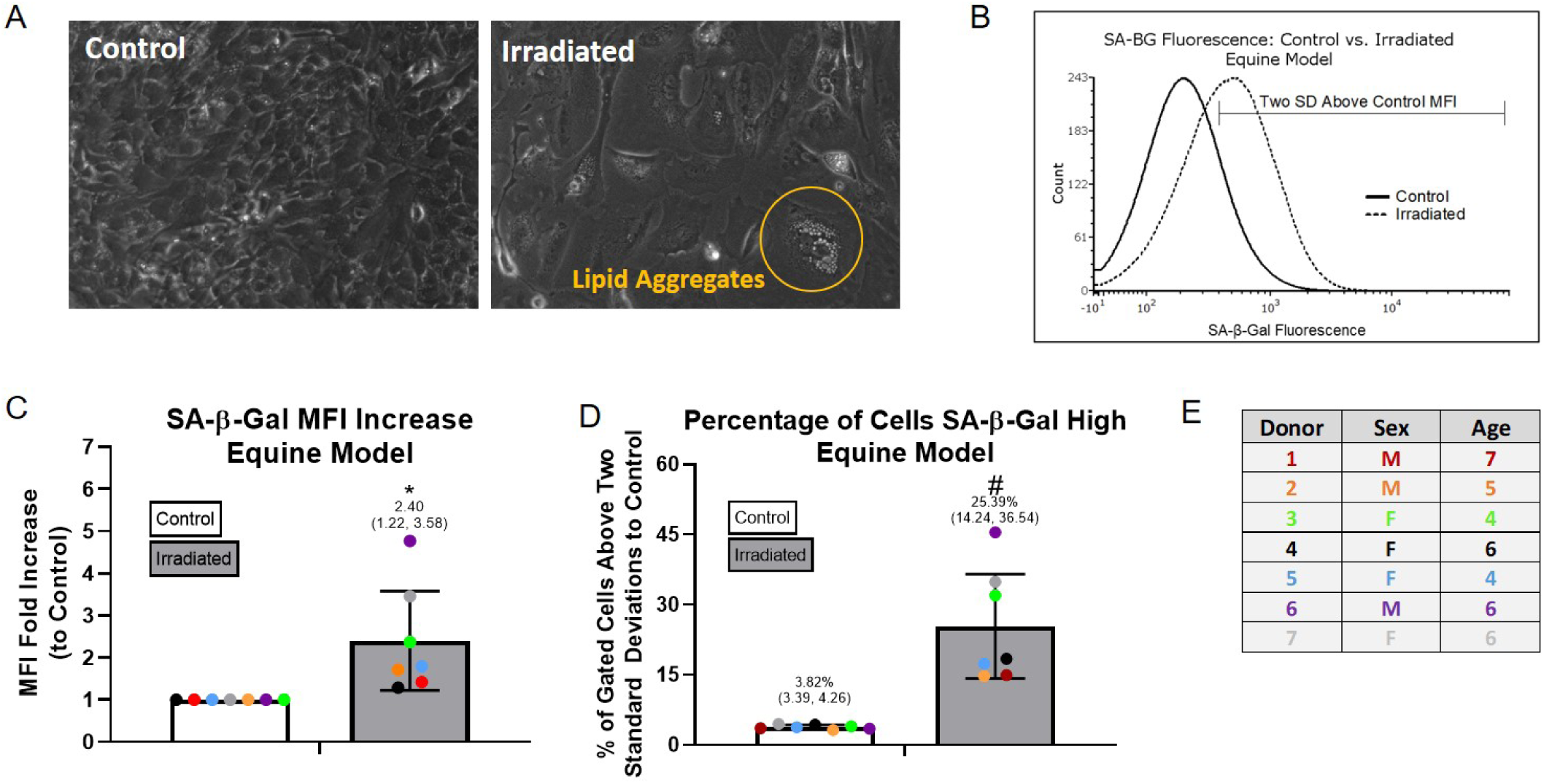
Quantification of senescence induction within equine explants. Equine explants were divided into control and irradiated groups, cultured in explant form for 7 to 10 days, digested and cultured in monolayer for an additional 10 to 12 days in 10% FBS for maturation of senescence phenotype. (A) Morphology of control and irradiated chondrocytes after 12 days of monolayer culture. (B) SA-β-Gal flow cytometry representative plot with the region two standard deviation above control MFI marked. (C) SA-β-Gal mean fluorescence intensity (MFI) fold increase above control quantified. The mean MFI fold increase from irradiated chondrocyte to control chondrocytes was 2.40; (*) indicates significance by one-sample t-test compared to a hypothetical value of 1, p=0.0272. (D) SA-β-Gal flow quantified as percentage of cells SA-β-Gal high. SA-β-Gal high cells delineated as those expressing SA-β-Gal fluorescence above 2 standard deviations from the control MFI. The mean percentage of SA-β-Gal high cells in the control group was 3.82%, whereas the mean percentage of SA-β-Gal high cells in the irradiated group was 25.39% (#) indicates significance by paired t-test, p=0.0031. (E) Information from equine donors included in the analysis (n=7 thoroughbred horses), no horses excluded from analysis.

### Induction of senescence in human cartilage explants

Preliminary studies on the human articular cartilage explants demonstrated a lack of senescence induction with irradiation alone as assessed by the SA-β-gal flow cytometry assay (data not shown). To stimulate a mitogenic response throughout the explant culture, we supplemented the 10% FBS media with growth factors TGF-β1 and bFGF during the senescence induction process. Cartilage explants harvested from cadaveric human ankles of 14 donors were cultured as described for the horse explants, with the only difference being the inclusion of growth factors during explant culture as an additional variable (**Fig. 1**). As compared to control chondrocytes, cells digested from irradiated explants cultured with growth factors showed an enlarged morphology and the presence of bi-nucleated cells, which has been associated with senescence^28^ (**Fig. 3a**). A representative SA-β-gal flow cytometry plot illustrates the robust shift in SA-β-gal fluorescence with both irradiation and growth factor treatment (**Fig. 3b**). Chondrocytes from irradiated explants treated with TGF-β1 and bFGF showed significantly higher SA-β-gal MFI values as compared to the non-irradiated 10% FBS control (1.44 mean fold increase, CI 1.26 to 1.63; p = 0.0002 by one-sample t-test compared to a hypothetical value of 1; **Fig. 3c**). By this measure, chondrocytes from irradiated explants without growth factors also showed a moderate increase in SA-β-gal MFI values as compared to the non-irradiated control (1.17 mean fold increase, CI 1.03 to 1.31; p = 0.025). When the percentage of SA-β-gal high cells was analyzed by Two-way ANOVA, both irradiation and media condition, as well as the interaction, were significant factors in senescence induction. Using Tukey’s multiple comparison test, the irradiated explants cultured with growth factors were significantly different than all other conditions (p<0.005 to all conditions) (**Fig. 3d**). A table listing the age and sex of the donors used in the irradiation experiments is provided (**Fig. 3e**).

**Figure 3.**
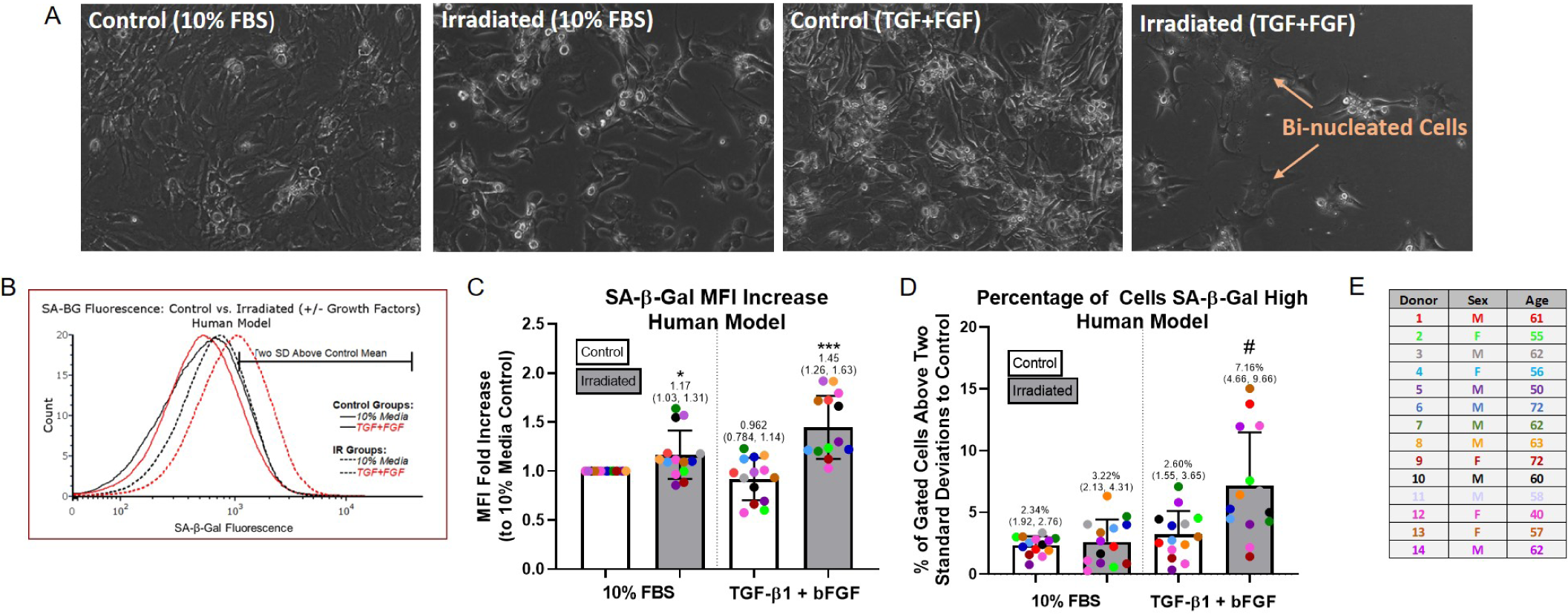
Quantification of senescence induction within human explants. Human explants were divided into control and irradiated groups, cultured in explant form for 7 days, then cultured in monolayer for an additional 12 days in 10% FBS for maturation of senescence phenotype. Growth factors, 1ng/mL TGF-β1 and 5ng/mL bFGF were added to the 10% FBS in the explant culture to provide an expansion stimulus. (A) Morphology of chondrocytes digested from control and irradiated cartilage (treated with or without growth factors) after 12 days of monolayer culture in 10% FBS medium. (B) SA-β-Gal flow cytometry representative plot with the region two standard deviation above control MFI marked. (C) SA-β-Gal MFI fold increase above control 10% condition quantified. Chondrocytes from irradiated explants treated with TGF-β1 and bFGF had a 1.45 mean MFI fold increase in SA-β-gal fluorescence; (***) indicates significance by one-sample t-test compared to a hypothetical value of 1, p=0.0002. Chondrocytes from irradiated explants treated in 10% FBS had a 1.17 mean MFI fold increase in SA-β-gal fluorescence; (*) indicates significance by one-sample t-test compared to a hypothetical value of 1, p=0.025.(D) SA-β-Gal flow quantified as the percentage of cells SA-β-Gal high. The mean percentage of SA-β-Gal high cells for the Control 10% FBS, Control TGF + FGF, IR 10%, and IR TGF + FGF conditions were 2.34%, 2.60%, 3.22%, and 7.16% respectively. (#) indicates significance to all other groups by 2-way ANOVA with Tukey’s multiple comparison test, p<0.005. (E) Information from human cartilage donors included in the analysis (n=14); three donors excluded due to too few cells (<800 events) in SA-β gal flow cytometry experiment and one outlier identified by Grubbs test and removed from subsequent analysis.

### Role of cell-cycle arrest in senescence induction

To explore the extent to which other forms of cell cycle arrest contribute to senescence induction, we used a pharmacological approach to avoid the extensive DNA damage caused by irradiation. Palbociclib inhibits the binding of CDK4/6 to D type cyclins^29^ and has been shown to completely inhibit the proliferation of human chondrocytes^30^. Explants derived from cadaveric human ankle cartilage were either treated with 1 µM of palbociclib for 12 days or treated with a vehicle control. Additionally, half of the explants in each group were treated with TGF-β1 and bFGF throughout the explant culture. As with the irradiation experiments, all groups received control medium in monolayer culture. Cadaveric human ankle cartilage from a total of 5 donors (distinct from the donors used in the irradiation experiments) were used for this study. A representative SA-β-gal flow cytometry plot illustrates a shift in SA-β-gal fluorescence for the explants treated with both palbociclib and growth factors (**Fig. 4a**). As compared to non-palbociclib treated 10% FBS control explants, SA-β-gal MFI increased in chondrocytes from explants treated with both palbociclib and growth factors (1.55 mean fold increase, CI 1.29 to 1.82; p = 0.0044 by one-sample t-test compared to a hypothetical value of 1; **Fig. 4b**). The combined treatment of palbociclib treatment and growth factors also significantly increased the percentage of SA-β-gal high cells (p<0.03 to all other conditions, Two-way ANOVA with Tukey’s multiple comparison test; **Fig. 4c**). A table listing the age and sex of the donors used in the palbociclib experiments is provided (**Fig. 4d**).

**Figure 4.**
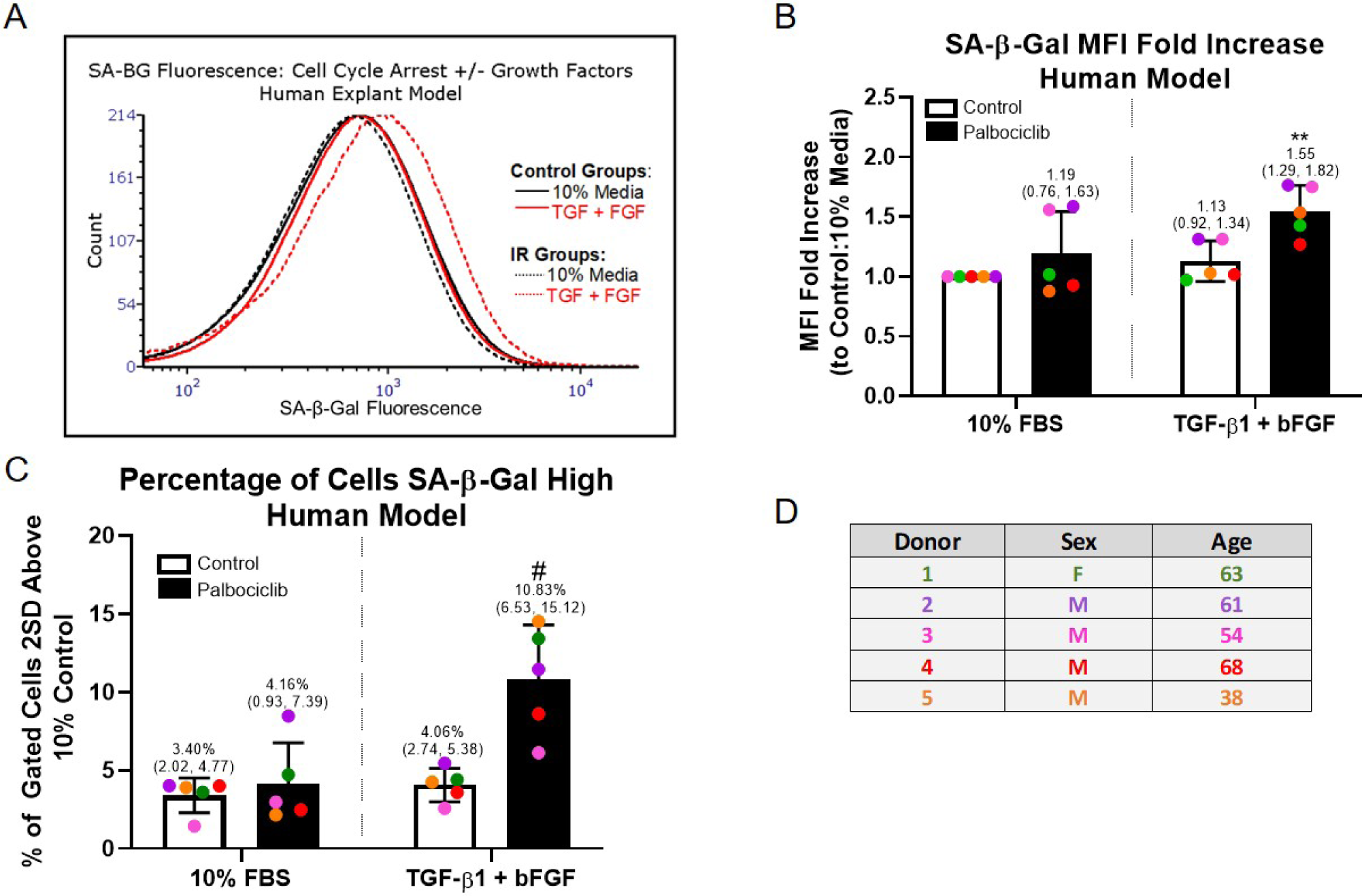
Role of cell-cycle arrest in senescence induction. An alternative method of stimulating cell-cycle arrest can be attained by treatment with 1μM of Palbociclib. Human explants were divided into control and Palbociclib treated groups, cultured in explant form for 7 days, then cultured in monolayer for an additional 12 days in 10% FBS for maturation of senescence phenotype. Growth factors were added to the 10% FBS media to provide an expansion signal. (A) SA-β-Gal flow cytometry representative plot of control and Palbociclib treated monolayer cells (+/-) the addition of TGF-β1 and bFGF. (B) SA-β-Gal MFI fold increase above control 10% FBS condition. The mean MFI fold increase of the Palbociclib condition with growth factors to the control 10% FBS group was 1.55; (**) indicates significance by one-sample t-test compared to a hypothetical value of 1, p<0.005. (C) SA-β-Gal flow quantified as the percentage of cells SA-β-Gal high. The mean percentage of SA-β-Gal high cells for the Control 10% FBS, Control TGF + FGF, Palbociclib 10%, and Palbociclib TGF + FGF conditions were 3.40%, 4.06%, 4.16%, and 10.83% respectively. (#) indicates significance to all other groups by 2-way ANOVA with Tukey’s multiple comparison test, p<0.05. (D) Table of the human cartilage donors included in the analysis, (n=5). One outlier identified by Grubb’s test and removed from analysis.

### Cluster formation in response to senescence-inducing conditions

H&E staining of cartilage at the end of explant culture revealed an increase in the presence of chondrocyte clusters of three or more cells in conditions that also resulted in senescence induction (irradiation and palbociclib). As with senescence induction, the inclusion of growth factors enhanced this response (**Fig. 5a**). There was a statistically significant increase in the percentage of chondrocyte clusters in a triplet^+^ formation for the palbociclib (plus growth factors) group as compared to the 10% FBS control group (p=0.015, Two-way ANOVA with Tukey’s multiple comparison test; **Fig. 5b**).

**Figure 5.**
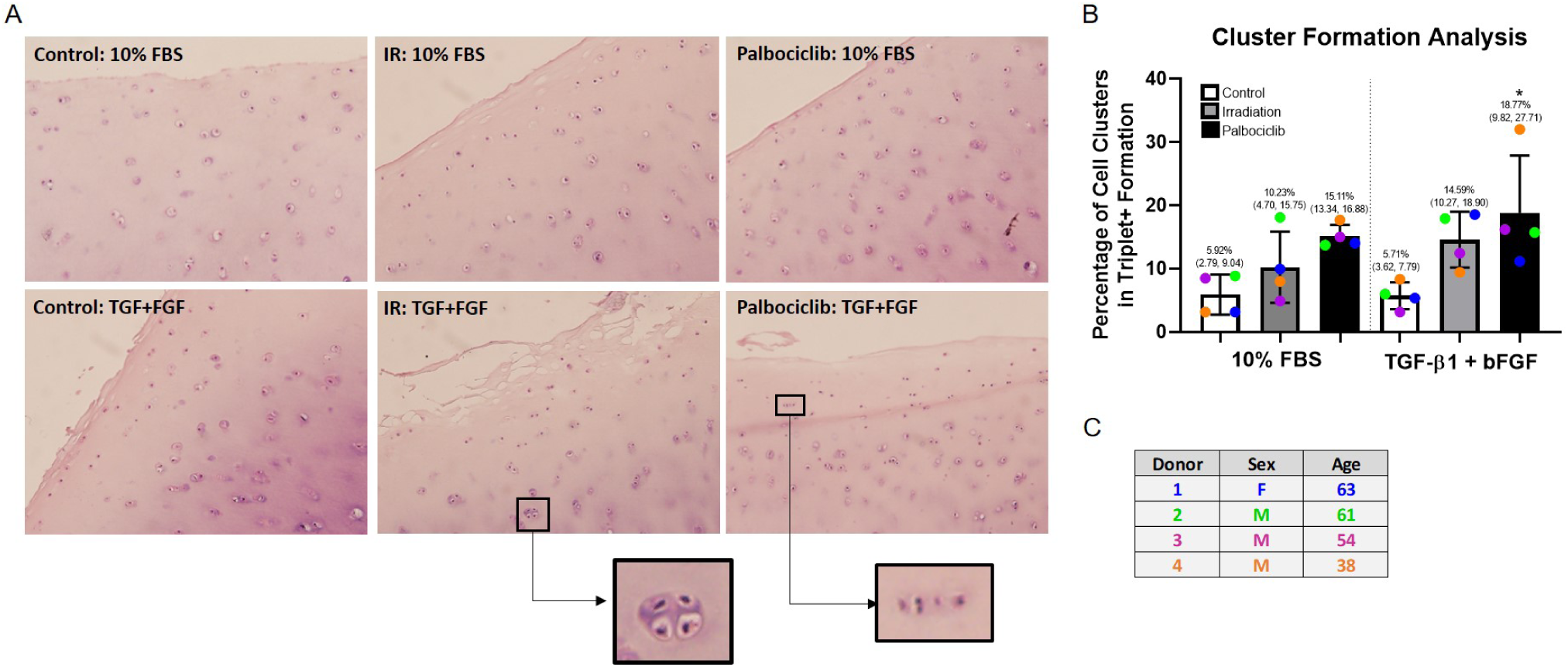
Chondrocyte clusters in cartilage explants correlate with senescence measures. 10% FBS media induced minimal proliferation within the human cartilage explants, indicating 10% FBS is not a strong enough stimulus to initiate chondrocyte division. Growth factors, 1ng/mL TGF and 5ng/mL bFGF were added to the 10% FBS to induce proliferation. (a) H&E staining on human explants after explant culture for 7 days, with or without growth factors. The presence of cell clusters in a triplet+ formation was more prevalent in the group receiving growth factors and a cell cycle arrest. (b) Cell cluster analysis on control explants (+/-) growth factors, irradiated explants (+/-) growth factors, and Palbociclib treated explants (+/-) growth factors. (*) indicates significance by 2-way ANOVA with Tukey’s multiple comparison test to the control groups, p<0.05 (c) Table of the human cartilage donors included in the analysis, (n=4). One outlier identified by Grubb’s test and removed from analysis.

## Discussion

This study demonstrates that treatment of articular cartilage explants with mitogenic stimuli in the context of cell-cycle arrest reliably induces high levels of SA-β gal activity (summarized in **Fig. 6**). By maintaining cell-matrix interactions, this approach provides a physiologically relevant setting to investigate the initiation of senescence. Detailed characterization of the origins of cellular senescence within cartilage may provide insight into why senescent cells accumulate to pathological levels in the joint space with age and in response to joint injury. Greater understanding of senescence induction will also support the emerging therapeutic approach to target senescent cells with senolytics^31,32^, which has shown promising results in animal models for post-traumatic and age-related OA^13^.

**Figure 6.**
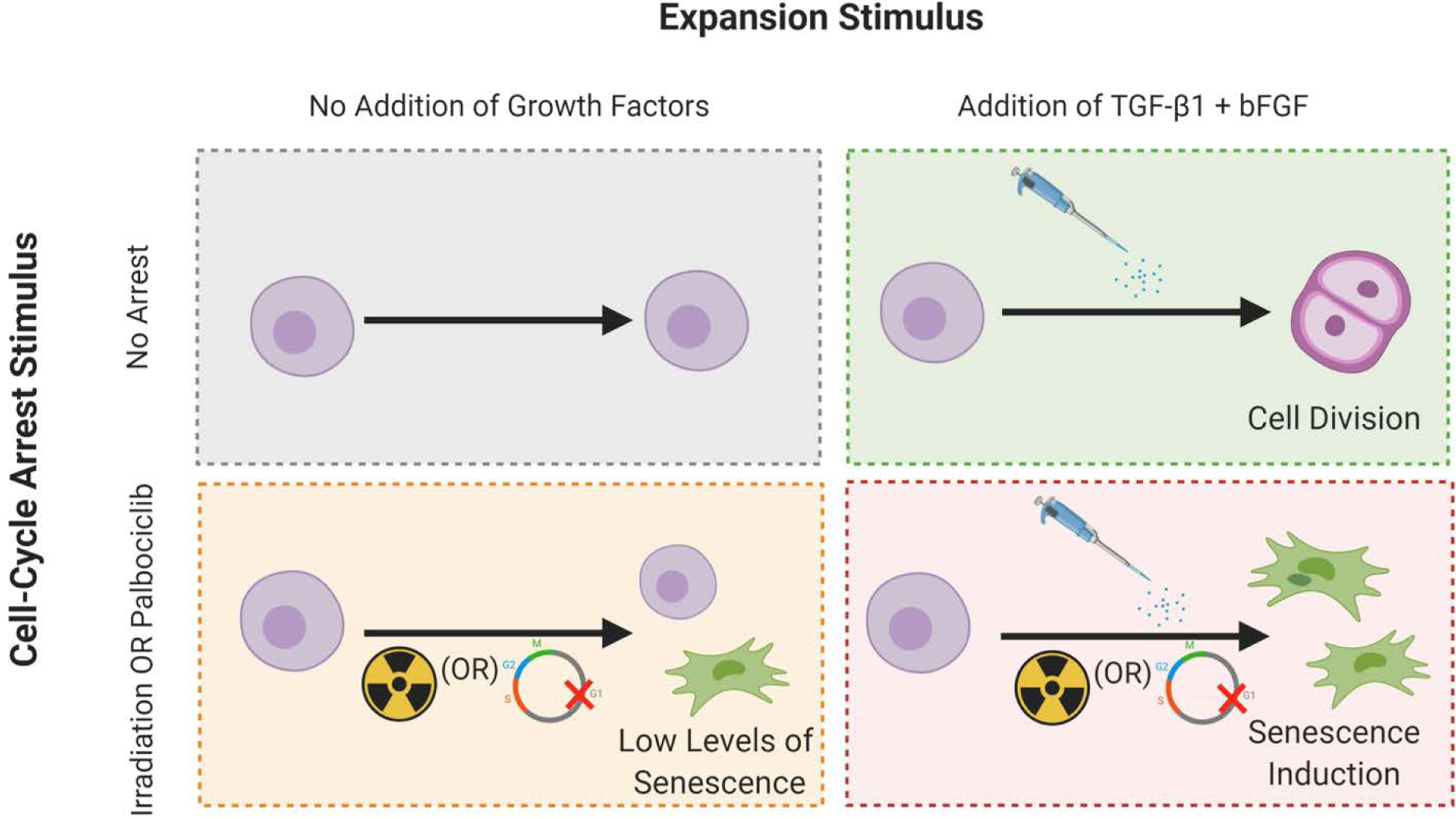
Overview of the senescence induction process. The senescence induction phenotype is hypothesized to be a result of conflicting cues from cell-cycle arrest stimuli and expansion stimuli. In the context of this study, the expansion stimulus was provided by 10% FBS (equine model) and 10% FBS + bFGF + TGF-β1 (human model), while the cell-cycle arrest stimulus was provided by either irradiation or palbociclib. Figure Created with BioRender.com.

The most widely used biomarker for senescence is SA-β-gal, which distinguishes senescent cells based on high lysosomal activity^33^. Traditionally, SA-β-gal is detected cytochemically using 5-bromo-4-chloro-3-indolyl-β-D-galactoside (X-gal) as a substrate and by discriminating negative from positive cells by visualization of blue color in cell culture images^34^. This manual process means that only a limited number of cells can be analyzed, and the readout is subjective and binary in nature. Conversely, the SA-β-gal flow cytometry approach utilized in this study provides a quantitative readout that captures the full range of lysosomal activity and is capable of analyzing tens of thousands of cells in a short time frame. Prior studies using a related but distinct flow cytometry readout of β-galactosidase activity revealed enhanced sensitivity as compared to the more widely used cytochemical method^34^. Although full characterization of the senescent state incorporates numerous orthogonal biomarkers^35^, the quantitative analysis of SA-β-gal was effective for distinguishing the cellular response to the applied culture conditions used in this study.

The cartilage utilized in this study originated from the equine stifle and from cadaveric human cartilage explants. The horse is a representative model of naturally occurring human osteoarthritis due to the similarities between human and equine articular cartilage and subchondral bone thickness^36^. Further, spontaneous joint injury is common in horses and post-traumatic OA is an important challenge in veterinary care^37,38^. High levels of senescence were induced in equine explants without the addition of growth factors, perhaps because the cartilage from relatively young horses showed a significant mitogenic response to the serum-containing medium. In addition to elevated SA-β-gal activity in the irradiated explant group, there was a notable change in chondrocyte morphology. This included the presence of lipid aggregates, which mirrors the “accumulation of lipids in senescence” phenotype previously described^27^. This phenotype suggests that senescent equine chondrocytes may be particularly susceptible to the reported senolytic role of fenofibrates^39^, as these drugs target lipid metabolism.

Cellular senescence was initially identified as an irreversible phenotype that cells entered upon reaching the limit of normal cell proliferation^40^. The observation that chondrocytes rarely replicate during cartilage homeostasis^41^ suggests that replicative senescence is not responsible for the accumulation of senescent cells in cartilage. However, the growth factors that are released from degrading matrix during early OA provide a potent mitogenic stimulus, as evidenced by the emergence of chondrocyte “clusters”. Increased numbers and sizes of cell clusters are characteristic of OA articular cartilage and these clusters are often localized in fissures and clefts of the upper cartilage layer^42–44^. bFGF is released from damaged cartilage tissue and has been identified near chondrocyte clusters^45^. There is also evidence of an interactive effect with TGF-β in cluster formation^46^, which is of particular interest given the finding that TGF-β is a key mediator of senescence in certain contexts^47^. In this study, human explants treated with TGF-β and bFGF showed high rates of senescence induction and cluster formation only when the tissue experienced conditions that restrain normal proliferation – irradiation or CDK4/6 inhibition. This finding is consistent with the concept that cellular senescence arises from the coordination of two conflicting processes – cell expansion and cell-cycle arrest^20,48–50^.

There are currently no cures for OA, and the development of effective treatments has been limited by an insufficient understanding of disease initiation and progression. This *in vitro* system establishes a set of physiological cues that induces senescence within both equine and human cartilage tissue, which will enable future mechanistic studies on senescence induction, the role of senescent chondrocytes in cartilage dysfunction, and the use of senolytic compounds to target senescent cells as a potential treatment for OA.

## Acknowledgements

The authors thank Mrs. Julie Long and members of the Central Procedures Laboratory at the North Carolina State University College of Veterinary Medicine for help in isolating equine cartilage explants as well as members of Dr. Richard Loeser’s laboratory at the University of North Carolina at Chapel Hill (UNC) for help in isolating human cartilage explants. The authors also would like to acknowledge the Gift of Hope Tissue and Organ Donor Bank and donor’s families as well as Dr. Arkady Margulis, MD, for tissue procurement. The authors appreciate assistance from the UNC Animal Histopathology and UNC Flow Cytometry Cores, the latter of which is supported in part by a P30 CA016086 Cancer Center Core Support Grant to the UNC Lineberger Comprehensive Cancer Center. Figures 1 and 6 of the manuscript were created using BioRender.com and exported under a paid subscription.

## Contributions

The conception and design of this study was done by MEC and BOD. Collection and analysis of data conducted by MEC, MCF, GAS and BOD. Study materials provided by RG, JMG, RFL, LVS, and SC. MEC drafted the article and RFL, LVS, SC, and BOD contributed to the critical revision of the article. All authors have read and approved the final manuscript.

